# Early, sex-dependent and progressive proteomic imbalance in the amygdala during Alzheimers disease progression

**DOI:** 10.1101/2025.11.26.690815

**Authors:** Silvia Romero-Murillo, Mercedes Lachén-Montes, Paz Cartas-Cejudo, Elena Anaya-Cubero, Marina de Miguel, Leire Extramiana, Isidro Ferrer, Joaquín Fernández-Irigoyen, Enrique Santamaría

**Affiliations:** Clinical Neuroproteomics Unit, Proteomics Platform, Sex & Gender in Health Research Area, Navarrabiomed, Hospitalario Universitario de Navarra (HUN), Universidad Pública de Navarra (UPNA), IdiSNA. Navarra Institute for Health Research, Pamplona, Spain; Department of Pathology and Experimental Therapeutics, UB, IDIBELL, CIBERNED-ISCIII, Barcelona, Spain

**Keywords:** Alzheimer’s disease, Amygdala, Proteomics, Sex differences, Drug repurposing

## Abstract

**Background:** The amygdala is involved in the emotional expression, memory processing and managing stimulatory input. Although amygdala atrophy is early evidenced in Alzheimer’s Disease (AD), the molecular mechanisms disrupted in initial neuropathological stages are still unknown. In the present study, we investigated the proteomic impairment of the amygdaloid region from AD-Braak stage I-II and III-IV subjects to better understand the neuropathological processes occurred early in this area and to identify potential targets that may face AD from the beginning of the disease.

**Methods:** Label-free quantitative proteomics was applied using an Orbitrap Exploris 480 mass-spectrometer in 24 postmortem amygdala specimens derived from non-demented (n=3F/5M), AD-Braak stage I-II (n=4F/4M) and AD-Braak stage III-IV (n=4F/4M). Data analysis was performed using MaxQuant and Perseus software (two-way Student T-test; p<0.05). Metascape and Ingenuity Pathway Analysis softwares were considered for biological interpretation. Connectivity map platform was used for drug repurposing analyses. Transcriptomic/proteomic data of other brain regions were obtained from AlzData, Neuropro, and Agora repositories.

**Results:** Amygdaloid proteome of AD-Braak stage I-II and III-IV subjects compared to controls revealed a progressive proteomic impairment with a minimal overlap across Braak stages. Some of the amygdaloid DEPs were known interactors of human Aβ plaques, APP, or Tau proteins or were previously identified at transcriptional or translational level in other brain regions affected by AD. Interestingly, amygdaloid proteome was more severely affected in women than in men with a particular protein expression profile associated to each AD stage. Comparing our sex-dependent differential proteome datasets with transcriptomic data of different brain regions, we identified potential sex-specific proteins related to cognitive decline and neurodegeneration. Finally, data-driven drug repositioning using amygdaloid omics profiles unveiled that most of the small molecule candidates were neuropathological stage and/or sex-specific.

**Conclusions:** Early and sex-specific amygdaloid proteome dysregulation in AD highlights the consideration of a deliberate stratification by sex in future research and clinical trials to develop effective therapeutic strategies in AD for both sexes.

**Plain English summary:** The amygdala is a brain region involved in the expression of emotions, memory processing and managing incoming stimulus. Atrophy of this area is evidenced at the first stages of Alzheimer’s Disease (AD), pointing out a potential involvement of amygdala in the pathology of this disease. However, the molecular changes occurred early in this area are not fully understood. To this end, we interrogated the proteome of amygdala postmortem samples came from subjects of early AD stages. By applying data and functional analyses, we observed a stage-dependent and progressive proteomic impairment in this area. We detected proteins differentially expressed that were already known to interact with well-stablished neuropathological proteins or were altered in other brain areas. Importantly, data stratification by sex revealed that protein expression changes of amygdala were more abundant in women than men across AD progression. After comparing our results with published data in different brain regions affected by AD, we identified sex-specific proteins that could be used as biomarkers of cognitive decline and neurodegeneration. Finally, a drug repositioning-based approach proposed candidates with the potential to reverse amygdaloid malignant AD signature more effectively in one sex than in other or just in one sex. These observations highlight the consideration to include sex differences in future research to develop more precise and effective treatments in AD.

**Highlights:** · Amygdaloid proteome experiences an increasing impairment across early neuropathological stages of AD, with a minimal overlap between Braak I-II and Braak III-IV

· Some of the amygdaloid DEPs are known interactors of human neuropathological Aβ plaques, APP, or Tau proteins, or are potentially connected with them direct or indirectly

· Protein expression changes in AD amygdala are more abundant in women than men across Braak staging, with a particular protein expression profile associated to each AD stage

· There are potential sex-specific proteins related to cognitive decline and neurodegeneration expressed in amygdala and other different brain regions

· Drug candidates that potentially reverse the neuropathological amygdaloid proteome are sex-specific, highlighting the need to consider sex stratification in future research to improve results translatability and progress on the field.

## Background

The amygdala is a central structure of the limbic system involved in three main functions: expression of emotions, memory processing and stimulatory input management. It maintains widespread connections with other brain regions, such as neocortical, allocortical and brainstem structures, which are thus affected by the dysregulation of each other [1].

In neurodegenerative diseases, the amygdala is significantly implicated in the disruption of brain system communication and behavioral output [1]. Pathologically misfolded proteins, such as neurofibrillary tangles of Tau and Aβ amyloid plaques, are often aggregated in the amygdala, especially in late Alzheimer’s disease (AD) [2].

Importantly, a pronounced amygdala atrophy has been shown in earlier stages of AD, positively correlated with symptom severity [3]. Also, preclinical studies using young APP/PS1 double-transgenic mice revealed reduced dendritic complexity of neurons, decreased levels of 5-HT, elevation of Aβ aggregates, and memory decline early in the amygdala [4,5]. Together, these findings on early studies hypothesize the amygdala as a key player in AD pathology and suggest the incorporation of its function into the screening for the diagnosis of early cognitive deterioration due to AD [1,5].

Advanced proteomics has emerged as a powerful approach to shedding light on the molecular mechanisms underlying the multifaceted AD pathology and identifying potential therapeutic targets [6,7]. Recent proteomic-based studies in AD have included different postmortem brain areas such as hippocampal and parahippocampal gyrus [8,9], frontal and temporal cortex [10], as well as olfactory-related regions [11–13], among others. However, proteomics analyses in amygdaloid complex are limited. Our group first characterized the proteome of healthy human amygdala, identifying 1814 unique proteins from which more than 60% of them had not been previously reported in the proteomes of other human limbic system structures [14]. Also, a study of nine brain areas from three individual brains thought to cover a spectrum of AD progression included amygdala sections in the proteomic analysis [15]. More recently, Gonzalez-Rodriguez et al combined stereological and proteomic analyses in non-Alzheimer’s disease and late AD amygdaloid human samples to describe synaptic alterations and potential participation of glial cells in response to pathology [16]. However, the molecular mechanisms underlying the neuropathological signatures shown in this area during initial AD stages are not fully understood.

In the present study, label-free quantitative proteomics and posterior bioinformatic and functional analyses of amygdaloid complexes from non-demented, AD-Braak stage I-II, and AD-Braak stage III-IV subjects reveal a progressive proteomic impairment with a minimal overlap as the disease progresses. Some of the amygdaloid DEPs were known interactors of human Aβ plaques, APP, or Tau proteins or were previously identified at transcriptional or translational level in other brain regions affected by AD. Considering sex dimension, the amygdaloid proteome was more severely affected in women than in men with a particular protein expression profile associated to each AD stage and sharing sex-specific co-expression patterns with other brain regions affected by AD, especially the parahippocampal and temporal cortex. Finally, data-driven drug repurposed candidates using amygdaloid omics profiles were sex-specific, highlighting the need to consider a deliberate stratification by sex in future clinical trials to develop effective therapeutic strategies in AD for both sexes.

## Material and methods

### Sample information and processing

#### Human samples

In accordance with Spanish Law 14/2007 of Biomedical Research, informed written consent for research purposes was obtained from the relatives of patients included in this study. Amygdaloid specimens and the corresponding clinical and neuropathological data from patients with Alzheimer’s disease (AD) were obtained from Institute of Neuropathology HUBICO-IDIBELL Biobank. The study followed the principles of the Declaration of Helsinki and all procedures received prior approval from the Clinical Ethics Committee of Navarra Health Service (study code: PI_2024/3). As previously described [11], left brain hemisphere was cut into 1-1.5 cm thick coronal sections, with the amygdaloid region and other brain areas rapidly dissected, frozen on metal plates over dry ice, and stored at -80°C for biochemical analysis. The other hemisphere was sunken in 4% buffered formalin for three weeks for morphological examination. Sections of the spinal cord were either frozen at -80°C or fixed in formalin for posterior analysis. Neuropathological diagnoses followed current guidelines, and neuropathological studies were conducted on paraffin-embedded sections from 25 brain regions, including the cerebellum, cerebrum, brainstem, and spinal cord, stained with hematoxylin and eosin, Klüver-Barrera, and periodic acid Schiff, or used for immunohistochemistry (antibodies against β-amyloid, phospho-tau (AT8), α-synuclein, αB-crystallin, TDP-43, TDP-43-P, ubiquitin, p62, glial fibrillary acidic protein, CD68, and IBA1). Cases with concurrent proteinopathies were removed from the study. For this study, 16 AD cases were grouped into Braak stage I-II (n = 4F/4M; mean age ± SD 69.1 ± 7.3), Braak III-IV (n = 4F/4M; mean age ± SD 79.1 ± 1.1) and eight additional specimens from elderly individuals without clinical history or histopathological evidence of neurological disease as controls (n = 3F/5M; mean age ± SD 66.1 ± 11.5) (**STable 1**).

### Amygdaloid proteomics

Protein extraction, proteome quantification, and analysis followed the workflow previously described [11].

#### Protein extraction, digestion, and purification

Human amygdaloid samples were lysed in buffer containing 7 M urea, 2 M thiourea, and 50 mM dithiothreitol (DTT), and centrifuged at 100,000 g for one hour at 15°C. Protein concentrations in the supernatants were measured using the Bradford assay (BioRad) and 20 μg of protein extracts per sample were reduced with 10 mM DTT during 30 minutes at room temperature (RT) and alkylated with 30 mM iodoacetamide for 30 minutes in darkness. An extra reduction step with DTT (30 mM) was carried out at RT for 30 minutes. The sample was then diluted to 0.6 M urea and trypsin (Promega; 1:50, w/w) was added for protein digestion at 37°C during 16 hours. Digestion was stopped by adding acetic acid and samples were purified using a vacuum manifold platform. Then, samples were dried, resuspended in 20 μL of 2% acetonitrile, 0.1% formic acid and 98% miliQ water, and quantified by NanoDropTM spectrophometer (Thermo).

#### Mass spectrometry

LC-MS/MS analysis was performed using an UPLC Ultimate-3000 system (Thermo) coupled to an Orbitrap Exploris 480 mass spectrometer (Thermo). 500 ng of peptides were separated using an analytical C18 Aurora column (75µm x 250 mm, 1.6 µm particles; Ionopticks) at a flow rate of 300 nL/min (40°C) using a 126-min gradient: 5% to 20% B in 100 min, 20% to 32% B in 25 min, and a final step of 32% to 95 B in 1 min (A= Formic acid 0.1%; B=100% Acetonitrile with 0,1% formic acid).

Mass spectrometer was operated in positive data-dependent acquisition mode. MS1 spectra were measured in the Orbitrap Exploris 480 mass spectrometer with a resolution of 120,000 (AGC target: Standard; maximum injection time: Auto; mass range: from 375 to 1,500 m/z; profile). The instrument was set to run in top speed mode with 3 sec cycles for the survey and the MS/MS scans. After survey scan, the most abundant precursors (with charge state between +2 and +5) were isolated in the quadrupole (Isolation window: 1.4 m/z) and fragmented in the Ion Routing Multipole HCD-Cell (collision energy: 30%). Fragmented precursors were detected in the Orbitrap (First mass: 110 m/z; AGC target for MS/MS: standard; maximum injection time: Auto; centroid). The dynamic exclusion was set to 30 sec with a 10 ppm mass tolerance around the precursor.

#### Data Analysis

All mass spectra were analyzed with MaxQuant software version 2.0.1.0. MS/MS spectra were searched against Homo sapiens Swissprot (isoforms) protein sequence database and GPM cRAP sequences (protein contaminants). Precursor mass tolerance was set to 20 ppm and 4.5ppm for the first search where initial mass recalibration was completed and for the main search, respectively. Product ions were searched with a mass tolerance 0.5 Da. The maximum precursor ion charge state used for searching was 5. Carbamidomethylation of cysteines was searched as a fixed modification, while methionine oxidation, acetylation of protein N-terminal, deamidation of glutamine/asparagine and pyro-Glu of glutamine were searched as variable modifications. Enzyme was set to trypsin in a specific mode and a maximum of two missed cleavages was allowed for searching. The target-decoy-based false discovery rate (FDR) filter for spectrum and protein identification was set to 1%.

The analysis of the Maxquant output file and subsequent visualization was done by Perseus software [17] following the protein identification and quantification criteria previously described [11]. After width-adjustment normalization, a 1.3-fold change cut-off was used for statistical significance (two-way Student T-test; p<0.05) and proteins with and absolute fold change below 0.77 were considered downregulated and those with higher range than 1.33 were considered upregulated.

#### Functional analysis and databases used

To identify the biological functions represented by the differentially expressed protein datasets, Metascape database [18] (http://metascape.org/) and QIAGEN’s Ingenuity Pathway Analysis (IPA) (version 107193442) (QIAGEN Inc; https://digitalinsights.qiagen.com/IPA) were employed. For Metascape analysis, default settings (minimum overlap: 3; minimum enrichment: 1.5; p<0.01) and Reactome database [19] were selected for enrichment analysis. IPA software calculates significant values between each biological event and the imported molecules based on Fisher’s exact test (p ≤ 0.05).

Connectivity Map (CMap) framework [20] was used for drug repurposing following the workflow previously applied in our group [21]. The differentially protein amygdaloid datasets obtained were uploaded to the CMap tool to identify drugs or genetic modifications that potentially reverse the proteomic fingerprint of each condition. Connectivity scores near to -100 express highly potential activity to reverse the proteomic signature.

Data from different brain regions of other AD-related omic studies were downloaded from Alzdata [22,23], Neuropro [24], or Agora [25] databases to compare common and specific patterns of expression between other brain areas and amygdala.

Figures were created with GraphPad Prism software v8.

## Results

### 1. Altered amygdaloid proteome in early neuropathological AD stages

To characterize the amygdaloid specific proteomic signature during AD progression, the proteomic profile of 24 amygdala post-mortem samples derived from neurologically intact controls and AD subjects in Braak I-II and III-IV stages was explored using label-free quantitative proteomics (**STable 1**). A progressive proteomic impairment was observed along the neuropathological progression. Sixty-six (3%) and 153 (6.9%) out of 2222 quantified proteins were differentially expressed in Braak I-II and Braak III-IV stages, respectively, compared to control group (FC 30%; p-value<0.05) (Fig. 1A**-B**; **STable 2-4**). Among them, only 13 (6%) differentially expressed proteins (DEPs) were commonly deregulated across Braak stages. TGM2, SHMT2, PRCP, TRIM3, NECAP1, PNPLA8, PDPK1, PDAP1, and BCAP31 were commonly downregulated, whereas ARPC5, PSMB6, ADIRF, and CLIP1 were upregulated (Fig. 1C). TRIM3, PNPLA8, and PDAP1 are known interactors of Tau protein, as well as BCAP31 with amyloid precursor protein (APP). In addition, NECAP1, PDAP1, PSMB6, SHMT2, TGM2, and TRIM3 have been previously detected in Aβ plaques (Fig. 1D) [26,27]. Moreover, a systems-biology approach allowed us to map direct and indirect functional relationships between Braak I-II or Braak III-IV amygdaloid DEPs and APP or MAPT (Tau) (Fig. 1E). Despite the modest proteome disruption at amygdaloid level, these results suggest that the alteration of the functional crosstalk between APP and Tau interactomes was more evident in intermediate Braak III-IV.

**Figure 1.**
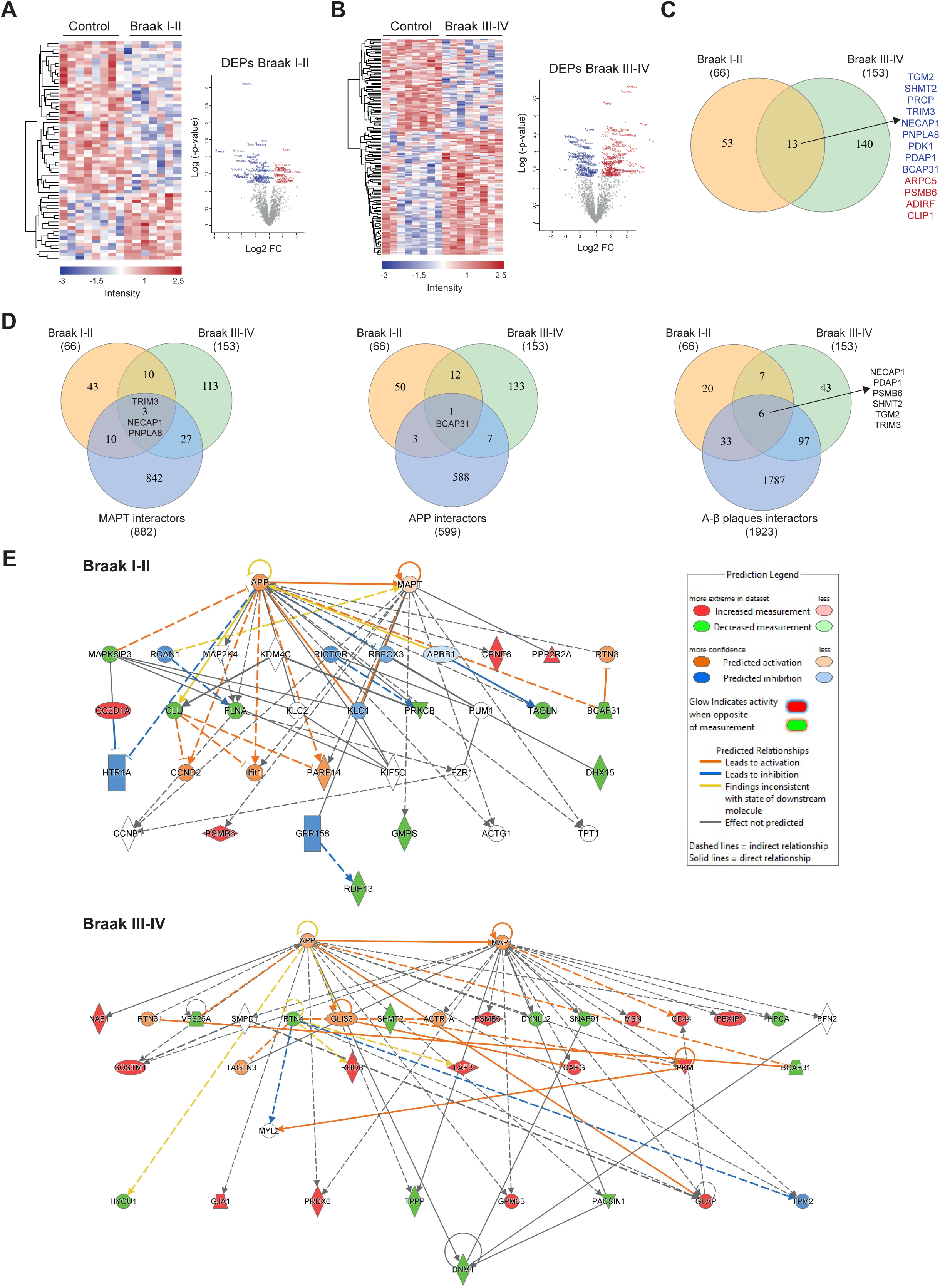
**Amygdala proteostatic imbalance during initial and intermediate AD stages**. Heatmap representation (left) and volcano-plots (right) of amygdala proteins differentially up-regulated (red) and down-regulated (blue) of AD subjects from (**A**) Braak I-II (n=7) and (**B**) Braak III-IV (n=8) stages compared to control samples (n = 8). (**C**) Venn diagram shows common altered proteins between Braak stages. Up-regulated and down-regulated common proteins are colored in red and blue, respectively. (**D**) Common proteins with known interactors of Tau (MAPT), APP, and human Aβ plaques. Figures created in <u>InteractiVenn.net</u> [59]. (**E**) Representation of the interactions between differentially expressed proteins in initial Braak I-II and Braak III-IV stages at amygdaloid level and APP and MAPT proteins according to IPA knowledgebase. DEPs, differentially expressed proteins.

In addition, we identified co-expression patterns in amygdaloid proteome with other brain regions during AD progression. Comparing our stage-dependent differential proteome datasets with published transcriptomic data of different brain regions of individuals with AD [22,23], the temporal cortex presented more similarities with amygdala, followed by entorhinal cortex, hippocampus, and frontal cortex (**S**Fig. 1A; **STable 5**). Consistent with the protein increased expression of ABLIM1 in amygdala samples of Braak I-II stage group, a significant upregulation of the gene encoded for this protein was observed in all four examined brain regions. Conversely, NECAP1 protein/gene exhibited a significant downregulation in all the comparison groups. PSMB6 was upregulated at protein level in amygdala whereas its gene levels were decreased in the rest of brain areas studied (**S**Fig. 1B; **STable 5**). Notably, 18 (11.8%) DEPs of intermediate Braak stages in amygdala (III-IV) displayed significant differential gene expression among all brain regions including *HPCA*, *RPH3A*, *NECAP1*, *GNG3*, *DNAJA2*, *APOO*, *SNAP91*, *GSS*, *SRM*, *PSMB6*, *ADD3*, *DTNA*, *GJA1*, *BBOX1*, *C12orf10*, *PBXIP1, GFAP*, and *CD44* (**SFig, 1C**; **STable 5**). When contrasted our results with 38 published AD proteomic studies from 13 different brain regions compiled in Neuropro database [24], the areas that matched more altered proteins in both stages were frontal (74 and 73% of Braak I-II and III-IV amygdaloid DEPs, respectively) and entorhinal cortex (35 and 50%) (**S**Fig. 1D; **STable 6**). Interestingly, part of the amygdaloid DEPs deregulated in other brain regions were also detected at transcriptional level when compared with AlzData database (**S**Fig. 1E; **STable 7**).

Despite the low protein overlap between stage-dependent proteotypes, functional enrichment analysis revealed common biological processes altered across staging including cellular responses to stress, carbohydrate metabolism, integrin-mediated signaling, signaling by Rho GTPases, axon guidance, neutrophil degranulation and clathrin-mediated endocytosis, pointing out selective metabolic trajectories that are initially modulated and maintained across intermediate Braak III-IV (Fig. 2A). However, protein intermediates associated to lipid, nucleotide and aminoacid metabolisms were specifically altered in Braak III-IV (Fig. 2A). The number of cellular functions aberrantly activated in this stage was greater, including mitotic-related processes, or KEAP1-NFE2L2 pathway, implicated in neuronal aging and AD neurodegeneration [28–30], among others. Interestingly, neutrophil degranulation pathway was predicted to be inhibited in Braak I-II and activated in Braak III-IV, being the most upregulated pathway predicted in this stage, suggesting an increase of the immune response and inflammation in intermediate III-IV stage. (Fig. 2B).

**Figure 2.**
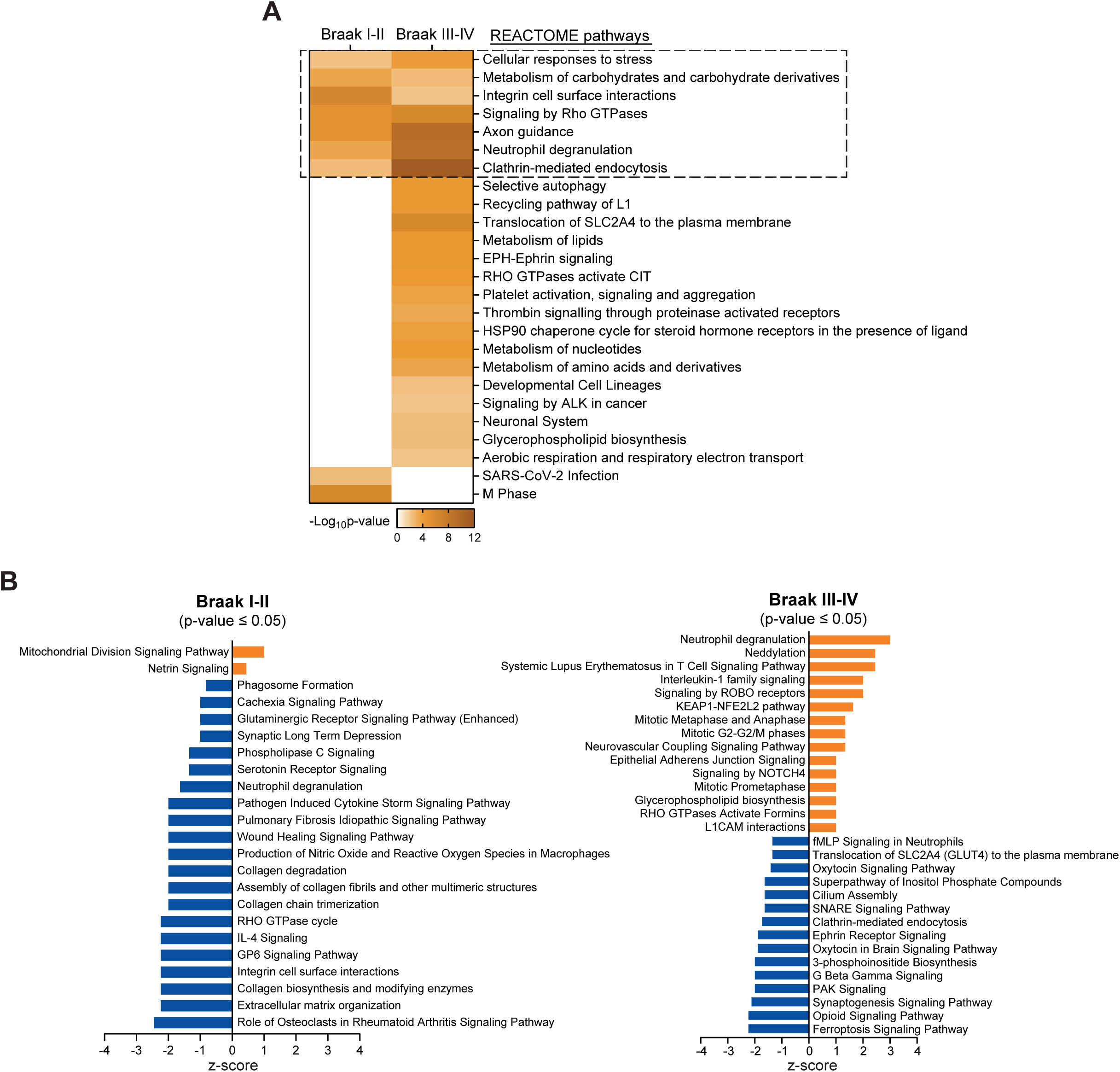
Functional enrichment analysis revealed common and specific biological processes altered across staging. (**A**) Functional mapping of disrupted amygdala proteome across AD neuropathological staging at pathway level using Reactome database [19]. Dashed lines indicate common pathways between both groups. (**B**) Prediction of activation/inactivation of canonical pathways in Braak I-II and Braak III-IV stages. Positive z-score values (orange) indicate potential up-regulated pathways whereas negative z-score values represent inhibited pathways with a p-value<0.05.

### 2. Sex-dependent amygdaloid proteome dysfunction in AD

Considering not only the neuropathological stage but also the sex dimension, the amygdaloid proteome was more severely affected in women than in men. Forty-four and 88 DEPs were identified in men across Braak staging (Fig. 3A**-B**; **STable 8-9**), whereas 73 and 137 DEPs were observed in women (Fig. 3A, D; **STable 10-11**), in accordance with the proteomic impairment described before. The overlap of DEPs between both sexes was minimal, sharing only 1 and 4 common proteins in Braak I-II and III-IV stages, respectively (**S**Fig. 2A**-B**). Moreover, 6 and 20 proteins were commonly deregulated across Braak staging in men and women, respectively (Fig. 3C, E). Interestingly, clustering analysis revealed different protein expression profiles across neuropathological grading between women and men. Although there were protein subsets with a progressive deregulated expression as the disease progresses (cluster 5 in men, cluster 1 in women), most of the differentially protein clusters were exclusively modulated in a neuropathological stage-dependent manner (Fig. 3F**-G**; **STable 12**). For instance, regarding men, specific protein subsets were downregulated (cluster 1) or upregulated (cluster 4) in Braak I-II stage, whereas clusters 2 and 3, with proteins involved in membrane trafficking and metabolism, respectively, were exclusively modulated later in the disease (Fig. 3F**, S**Fig. 2C). In women, cluster 2 and 3 were composed of proteins with increased expression only in Braak I-II stage.

**Figure 3.**
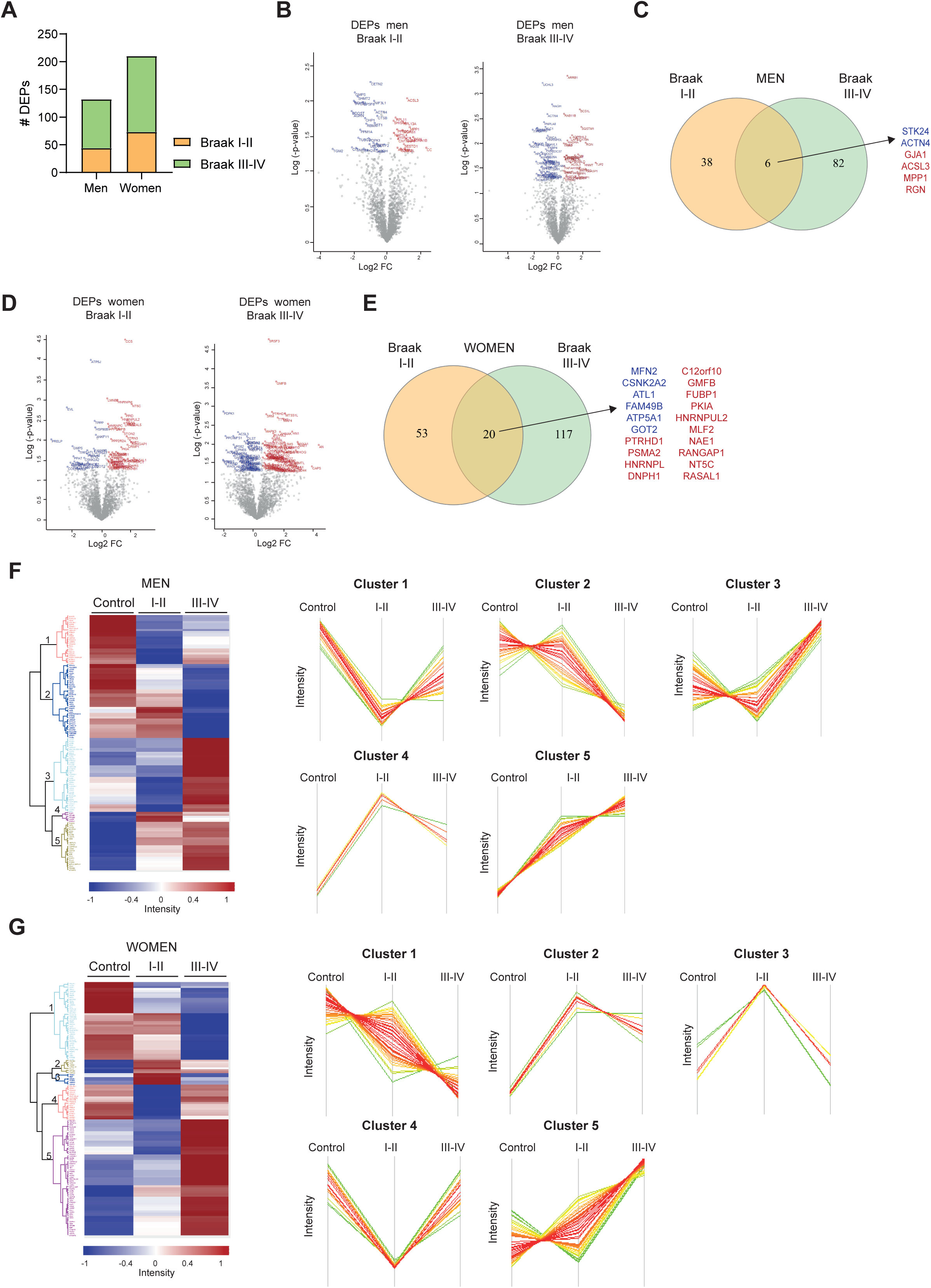
Amygdaloid proteome dysfunction is more severe in AD women than men. (**A**) Bar graph represents the number of DEPs of AD amygdala proteome in women and men across Braak staging compared with control groups. Volcano plots indicate the amygdala proteins significantly up-regulated (red) and down-regulated (blue) in Braak I-II and Braak III-IV stages in (**B**) men and (**D**) women subjects. Number of common and dissimilar deregulated proteins between Braak stages in (**C**) men and (**E**) women. Up-regulated and down-regulated common proteins are colored in red and blue, respectively. Figure created in <u>InteractiVenn.net</u> [59]. Heatmap representing the differential amygdala proteotyping in AD across Braak stages in (**F**) men and (**G**) women. Cluster plots show protein groups exclusively modulated in one Braak stage or with a progressive dysfunction across the disease. DEPs, differentially expressed proteins.

Proteins from cluster 4 showed a drop that recovered control levels in Braak III-IV, whereas proteins that belong to cluster 5 demonstrate an up-regulated expression (Fig. 3G**, S**Fig. 2C). Altogether, these observations support a specific and sex-dependent protein alteration pattern in each neuropathological AD stage.

At functional level, the overlap of altered pathways between both sex and Braak stages was modest, with nervous system development and signaling by Rho GTPases commonly imbalanced across all conditions (Fig. 4A). As expected, there were more cellular functions altered in women Braak stage III-IV. In particular, oxidative phosphorylation, complex I biogenesis, TCA cycle, and other mitochondrial metabolism-related pathways were predicted to be inhibited in this group. On the other hand, mitotic and immune functions tent to be activated (Fig. 4B). Regarding men, inositol phosphatase associated pathways were potentially downregulated in late Braak stages (Fig. 4B). As a coordinator of metabolic response including growth factor signaling, nutrient sensing, and cellular energy storage homeostasis [31], the inhibition of this system may indicate cell metabolism dysfunction. These results suggest that metabolism deregulation is commonly affected at Braak III-IV stage regardless of sex.

**Figure 4.**
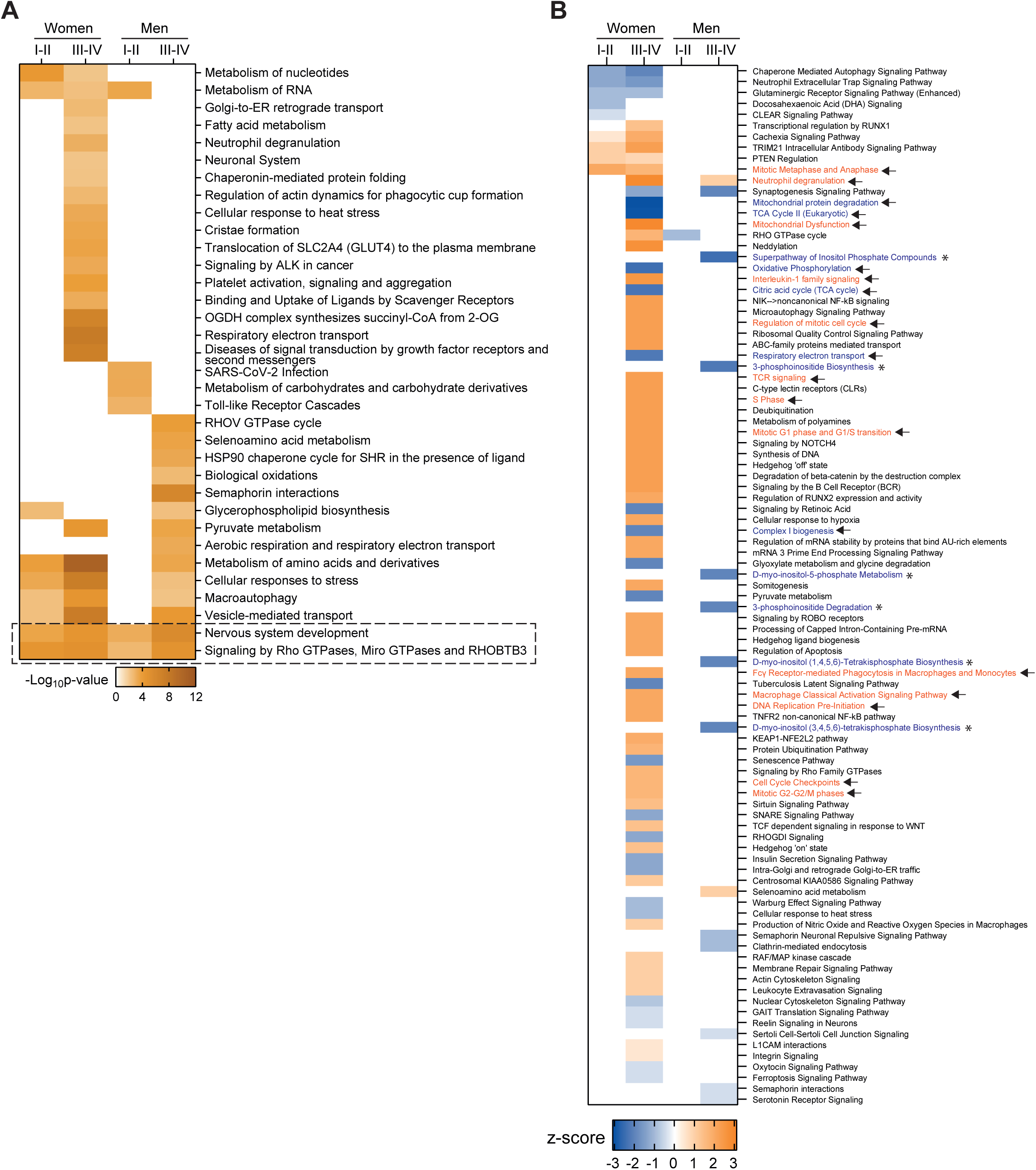
Sex-specific biological alterations across Braak Staging. (**A**) Functional mapping of disrupted amygdaloid proteome in AD women and men across Braak staging using Reactome database [19]. Dashed lines indicate common pathways between both groups. (**B**) Heatmap comparing the prediction of activation/inactivation of canonical pathways in AD women and men across Braak stages. Positive z-score values (orange) indicate potential up-regulated pathways whereas negative z-score values represent inhibited pathways (p-value<0.05). Arrows mark the pathways cited in the text.

### 3. Amygdaloid proteome dysfunction shares sex-specific co-expression patterns with other brain regions affected by AD

We contrasted our sex-dependent differential proteome datasets with transcriptomic data of 9 different post-mortem brain regions from over 1100 individuals of 3 human cohort studies grouped in Agora database [25]. Importantly, the data of this repository show relative changes in gene expression between female or males AD patients and their sex-respective control groups, allowing us to identify sex-dependent protein expression patterns in amygdala and other brain regions related to neurodegeneration in AD.

Compared to Agora database, parahippocampal gyrus (PHG) and temporal cortex (TCX) were the brain areas that presented more similarities with all amygdaloid datasets, with 14 and 20 (men Braak I-II), 40 and 26 (men Braak III-IV), 30 and 29 (women Braak I-II), and 68 and 58 (women Braak III-IV) protein-coding genes commonly altered with PHG and TCX, respectively (Fig. 5A). Among the DEPs obtained in men I-II Braak dataset, increased Gap Junction Alpha-1 protein (GJA1) and downregulated CHP1 phosphoprotein were identified in 6 of the 9 brain regions studied (Fig. 5B); whereas DPYSL3, involved in response to axon injury, and RPH3A were the proteins with more incidence from this analysis in Braak III-IV stages (men) (Fig. 5C). Regarding early AD stage DEPs in women’s amygdala (Braak I-II), HDGF and PRELP were also deregulated at transcriptional level in 8 out of 9 female brain areas affected by AD, followed by the guidance receptor Plexin-B1 (PLEKHB1), which regulates peri-plaque glial net activation in AD [32], TCEAL5 and ARF5 (Fig. 5D). In relation to amygdaloid women Braak III-IV proteome, glutathione synthetase (GSS) was detected in all of the brain regions from Agora database, although its gene expression did not correlate with the increased protein levels identified in AD women amygdala. Also, HPRT1 or AHNAK were also imbalanced in multiple female AD-affected regions (Fig. 5E).

**Figure 5.**
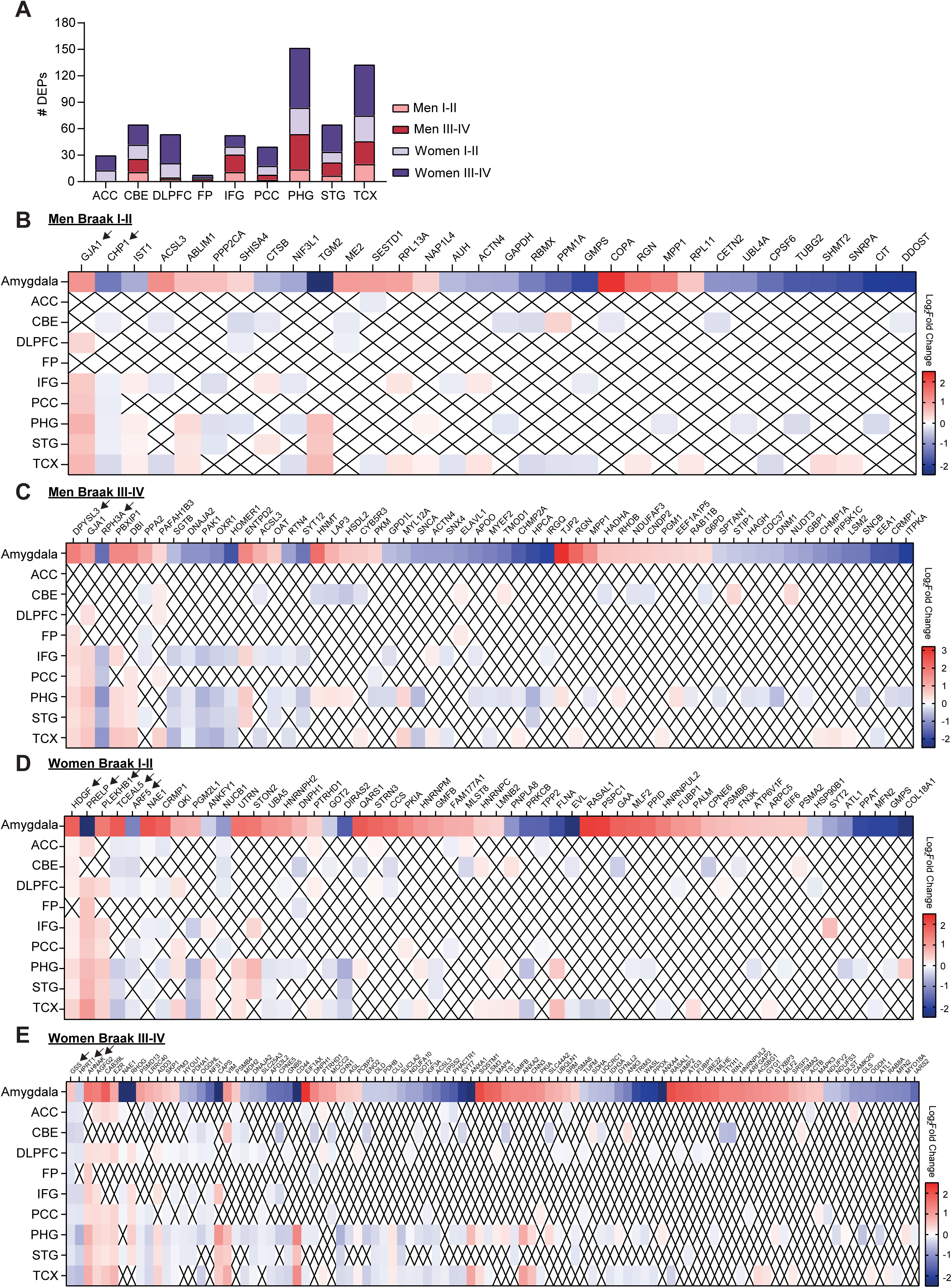
Amygdaloid proteome dysfunction shares sex-specific co-expression patterns with other brain regions affected by AD. (**A**) Number of amygdaloid DEPs from Braak I-II and III-IV datasets in men and women whose mRNA expression levels were also deregulated in other male or female brain regions affected by AD published in Agora database [25]. Heatmaps representing Log_2_Fold change of the proteins/genes differentially expressed in (**B**) men Braak I-II, (**C**) men Braak III-IV, (**D**) women Braak I-II, and (**E**) women Braak III-IV amygdala samples and other brain regions studied. Arrows mark the protein-coding genes cited in the text. DEPs, differentially expressed proteins; ACC, Anterior cingulate cortex; CBE, Cerebellum; DLPFC, Dorsolateral prefrontal cortex; FP, Frontal pole; IFG, Inferior frontal gyrus; PCC, Posterior cingulate cortex; PHG, Parahippocampal gyrus; STG, Superior temporal gyrus; TCX, Temporal cortex.

Moreover, Agora database calculates a Target Risk Score (TRS) to represent the gene target’s general relevance to AD based on genetic and transcriptomic/proteomic evidences (from 0 to 5, higher scores indicate a greater likelihood of disease association). On this matter, AGRN (TRS: 4.33), critical in the development of the neuromuscular junction, and MFN2 (TRS: 4.12), related to mitochondrial fusion, were the protein-coding genes with the highest TRS from Braak I-II amygdaloid datasets from men and women, respectively, whereas SQSTM1, that functions as a scaffolding/adaptor protein to mediate activation of NF-kB signaling pathway [33], was the top one in Braak III-IV in both sexes, with a TRS of 4.5 (**STable 13**).

Taken together, these analyses suggest potential sex-specific proteins related to cognitive decline and neurodegeneration that could be used as therapeutic targets in future studies considering both the neuropathological stage and the sex dimension for better clinical outcomes.

### 4. Drug repositioning based on the reversion of amygdaloid omic AD signatures

Following the drug repurposing workflow previously applied in our group [21], we used the CMap framework [20] to identify drugs or genetic modifications that potentially reverse or mimic the AD phenotype showed in each AD Braak stage and distinguishing by sex at the level of the amygdala. Nicotinamide, which is being tested in a phase 2 clinical trial in early AD (NCT03061474) [34], nornicotine, vindesine, or the PAR1 antagonist SCH-9797 were the leading compounds (highly negative connectivity scores) that could reverse the proteomic signature of both Braak I-II and Braak III-IV stages (Fig. 6A). Moreover, the overexpression of the tumor suppressor TSSC4 or the inhibition of protein-coding genes such as CLIC4, COG7, RAD51, or the heat shock proteins HSPA5 and HSPA8 were aimed to restore the metabolic imbalance from initial AD stages in amygdala (Fig. 6B**-C**; **STable 14**).

**Figure 6.**
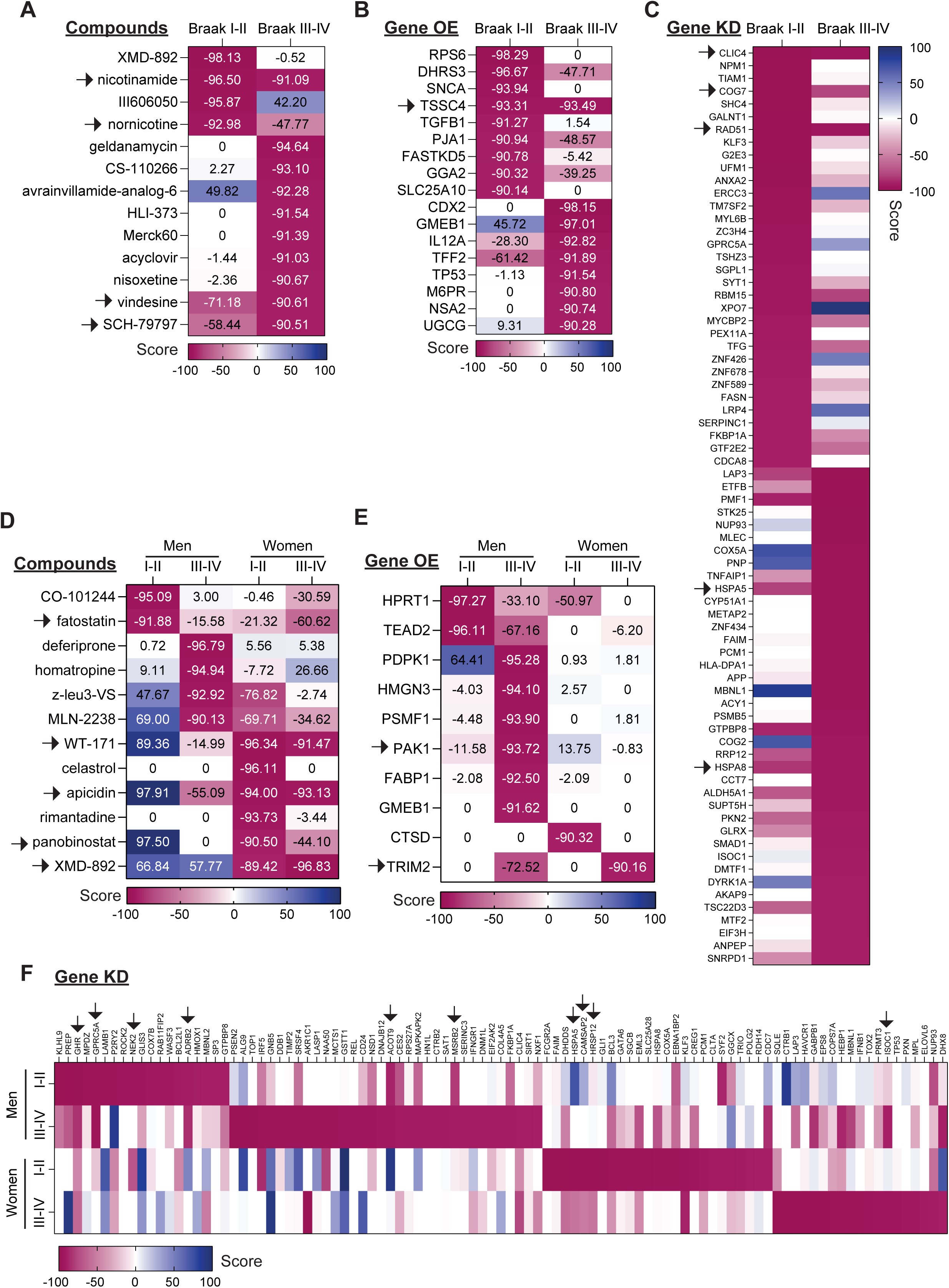
Potential candidates to reverse the amygdaloid proteomic AD signature. Heatmap showing top (**A**) potential pharmacological candidates and (**B**) genes whose overexpression or (**C**) genetic silencing reverse the proteomic signature of amygdala DEP datasets from each Braak stage and (**D, E, F**) considering sex. Connectivity scores near to -100 express highly potential activity to reverse the proteomic signature. Results were obtained from connectivity map database [20]. Arrows mark the compounds/genes cited in the text.

Interestingly, when considering not only the AD stage but also the sex dimension, the potential drugs and genetic modifications proposed by this workflow were quite different. Fatostatin, which inhibits the activation of SREBP, was the unique compound that potentially could reverse the omic phenotype of Braak I-II and Braak III-IV in both sexes, whereas HDAC inhibitors (WT-171, apicidin, or panobinostat) seemed to be effective only in women (Fig. 6D). Importantly, while XMD-892, a selective dual inhibitor of ERK5 kinase and bromodomain-containing proteins, was the top candidate to recover Braak I-II amygdaloid deregulated proteome (Fig. 6A), this drug possessed positive connectivity scores in both AD stages from men group (Fig. 6D). In the same way, most of the gene candidates whose overexpression or silencing may reverse the proteomic signature were specific of AD stage and/or sex. Stabilization of TRIM2, described to have a neuroprotective role [35], could be efficacious to oppose the amygdaloid omic fingerprint of Braak III-IV in both sexes (Fig. 6E). Interestingly, PAK1, which was downregulated in men amygdaloid proteome from Braak III-IV (**STable 9**) and in other four male AD-affected brain regions (Fig. 5C), had a connectivity score of - 93.72 to potentially reverse the proteomic signature of this group by its overexpression (Fig. 6E). Regarding the potential benefit of inhibiting genes, NEK2 and ISOC1 seemed to be effective in Braak I-II and Braak III-IV stages, respectively, regardless of the sex. On the other hand, G protein-coupling receptor family members (GPRG5A and ADRB2), ACOT9, and MSRB2 genes displayed negative connectivity scores in men, whereas HSPA5, HRSP12, and CAMSAP2 were significantly negative in women. Only the growth hormone receptor GHR showed negative connectivity scores in all for groups (Fig. 6F; **STable 15**).

## Discussion

Neuropathological accumulation of Aβ amyloid peptides and neurofibrillary tangles of hyperphosphorylated Tau are usually seen in amygdala on advanced AD [2]. Moreover, early amygdala atrophy, together with other clinical evidence of neurodegeneration in this area [3–5], suggest a significant implication of amygdala from the beginning of the disease. However, the molecular mechanisms disrupted in this region during initial neuropathological stages of AD are still unknown.

To this end, we investigated the proteomic impairment of amygdaloid complexes from AD-Braak stage I-II and III-IV subjects in the present study. We found 66 and 153 DEPs in Braak I-II and Braak III-IV stages, respectively, with a minimal overlap between them, indicating a stage-specific proteome disruption in amygdala. In accordance with the progressive proteomic impairment described in this work, previous proteomic data analysis of amygdaloid samples from more advanced stage (Braak V-VI) identified 178 DEPs in this group [16]. When comparing these data with our amygdaloid Braak I-II and Braak III-IV datasets (data not shown), only ADIRF protein seemed to be commonly upregulated across Braak staging, while the overlap between biological pathways was greater, with a progressive deregulation of cellular response to stress or neutrophil degranulation as the disease progresses.

By comparing amygdaloid DEPs with lists of known interactors of human Aβ plaques, APP, or Tau proteins, or mapping direct and indirect potential relationships with these neuropathological proteins, we identified amygdaloid proteins altered early in AD that could be directly implicated in the progressive depositions and aggregations of Aβ and NFT of Tau later in the disease. Moreover, after contrasting our results with transcriptomic and proteomic data of different brain regions affected by AD, we found that temporal, frontal, and entorhinal cortex presented more common dysregulated gene/proteins with amygdala than other brain areas. Even though, overall protein expression pattern pointed to be predominantly region specific. These results strengthen the need for broaden these studies to more brain areas for a better understanding of AD etiology and region vulnerability to disease [36].

The prevalence and incidence of AD differ by sex, being almost twice higher in women than men [37]. Therefore, understanding the mechanisms underlying these sex differences in AD is crucial to design effective treatments for both sexes [38,39]. A study based on postmortem data on nearly 1500 women and men showed that women had higher levels of AD pathology and tau tangle density irrespective of age at death compared to men [40]. Sex chromosomes and hormones, such as high FSH or low testosterone, contribute to AD pathological proteins deposition and impaired cognition [38,41–44]. Also, recent multi-omic studies reveal sex-specific molecular networks, differences in microglial signature and miRNA regulation, and metabolic and proteomic variations that may drive AD pathology differently in men versus women [38,45–48]. Here we showed that protein expression changes of amygdala were more abundant in women than men across Braak staging, with a particular protein expression profile associated to each AD stage. In accordance with our results, we previously observed a more severe dyshomeostasis of women olfactory bulbs affected by AD or Parkinson’s disease [11]. Importantly, the overlap between both DEPs and dysregulated biological pathways was minimal, pointing out a sex-specific impaired proteostasis of amygdala in AD. In both sexes, cellular metabolism seems to be affected at Braak III-IV stage. However, functional analyses suggested a direct mitochondrial metabolic impairment in women and a dysregulation of inositol phosphatase-related metabolic coordination in men.

Comparing our sex-dependent differential proteome datasets with transcriptomic data of 9 different brain regions from Agora database [25], we identified potential sex-specific proteins related to cognitive decline and neurodegeneration. For instance, GJA1, a regulator of microglial reactivity in AD [49] or RPH3A, correlated with dementia severity, cholinergic deafferentation, and increased Aβ concentrations [50] were imbalanced in multiple male AD-affected regions; whereas high levels of HDGF, or AHNAK, which has been recently identified as a key driver in the protein network disrupted in AD [8], were detected in female AD-affected areas.

Given the unmet clinical need for disease-modifying treatments in AD, drug repurposing has become an attractive approach due to its advantages in development timelines and costs [51]. Indeed, repurposed agents account for one-third of the AD pipeline agents in 2025 [52]. Through *in silico* evaluations, computational resources propose promising candidates to test in *in vitro* and *in vivo* settings [53]. In the present study, by levering our stablished drug repurposing workflow using the CMap framework [20,21], we identified drugs or gene modifications with the potential to reverse amygdaloid AD omic fingerprint in a sex-dependent manner. Interestingly, HDAC inhibitors were predicted to counteract AD amygdaloid proteome only in women, which may be explain due to their direct implication in the regulation of female hormones receptors [54]. These and other drugs, such as XMD-892, or gene modifications were proposed to have better response in one sex than in other or even be effective only in one sex. Therefore, ignoring these sex differences could difficult research results translatability and, thus, progress on the field. As an example, lecanemab, a recently approved anti-Aβ therapy for AD, demonstrated much more limited clinical benefit in women than men [55].

## Perspectives and Significance

Our data in AD amygdaloid samples from Braak I-II and Braak III-IV subjects contribute to the repertoire of the human brain deregulated proteome in AD and provide novel insights into the molecular mechanisms dysregulated early in this area. Sex differences in amygdaloid proteome support the significant impact of sex on prevalence, mechanism of disease and treatment response observed in AD patients [37,38,55]. Hence, it is crucial to include sex differences into both *in vitro* and *in vivo* models to develop more precise and translational research approaches for AD investigation [56].

## Conclusions

Amygdala proteome is dysregulated in early AD stages, supporting an implication of this brain area in AD pathology and progression from the beginning of the disease. Importantly, sex-specific protein expression patterns and therapeutic agents highlight the consideration of a deliberate stratification by sex in future researches and clinical trials to develop effective therapeutic strategies in AD for both sexes.

## Declarations

### Ethics approval and consent to participate

According to the Declaration of Helsinki, all assessments, postmortem evaluations, and experimental procedures were previously approved by the Clinical Ethics Committee of Navarra Health Service (study code: PI_2024/3). According to the Spanish Law 14/2007 of Biomedical Research, informed written consent from relatives of subjects included in this study.

### Consent for publication

All authors gave final approval of the manuscript and are accountable for all aspects of the work.

### Availability of data and materials

The mass spectrometry proteomics data have been deposited to the ProteomeXchange Consortium (http://proteomecentral.proteomexchange.org) via the PRIDE partner repository [57] with the dataset identifier PXD069452 (Access key for reviewers: BNhXUtL9sf17). According to recent recommendations [58], sex annotation has been included in raw files to facilitate further analysis.

### Competing interests

The authors declare no conflicts of interest.

### Funding

The Clinical Neuroproteomics Unit is supported by grants PID2023-152593OB-I00 funded by MCIU/AEI/ 10.13039/501100011033 / FEDER, UE to ES and JFI and 0011-1411-2023-000028 (from Government of Navarra- Department of Economic and Business Development-S4) to ES. Paz Cartas-Cejudo is supported by a postdoctoral fellowship from Public University of Navarra (UPNA).

### Authors’ contributions

Conceptualization: Enrique Santamaría; sample collection & neuropathology: Isidro Ferrer; data curation: Silvia Romero-Murillo, Enrique Santamaría; formal analysis: Silvia Romero-Murillo, Mercedes Lachén-Montes, Paz Cartas-Cejudo, Elena Anaya-Cubero, Marina de Miguel, Leire Extramiana, Joaquín Fernández-Irigoyen, Enrique Santamaría; funding acquisition: Enrique Santamaría, Joaquín Fernández-Irigoyen; investigation: Silvia Romero-Murillo, Mercedes Lachén-Montes, Paz Cartas-Cejudo, Elena Anaya-Cubero, Marina de Miguel, Leire Extramiana, Isidro Ferrer, Joaquín Fernández-Irigoyen, Enrique Santamaría; methodology: Silvia Romero-Murillo, Mercedes Lachén-Montes, Paz Cartas-Cejudo, Elena Anaya-Cubero, Marina de Miguel, Leire Extramiana, Isidro Ferrer, Joaquín Fernández-Irigoyen, Enrique Santamaría; writing—original draft: Silvia Romero-Murillo and Enrique Santamaría.

## Acknowledgements

We are very grateful to the patients and relatives who generously donor the brain tissue for research purposes. We are indebted to the Institute of Neuropathology HUBICO-IDIBELL Biobank for providing us the amygdaloid specimens as well as the associated clinico-pathological data. Authors thank the PRIDE Team for helping with the mass spectrometric data deposit in ProteomeXChange/PRIDE. The Clinical Neuroproteomics Unit of Navarrabiomed is a member of the Spanish Olfactory Network (ROE) (supported by grant RED2022-134081-T funded by Spanish Ministry of Science and Innovation).

---

**Supplemental figure 1.**
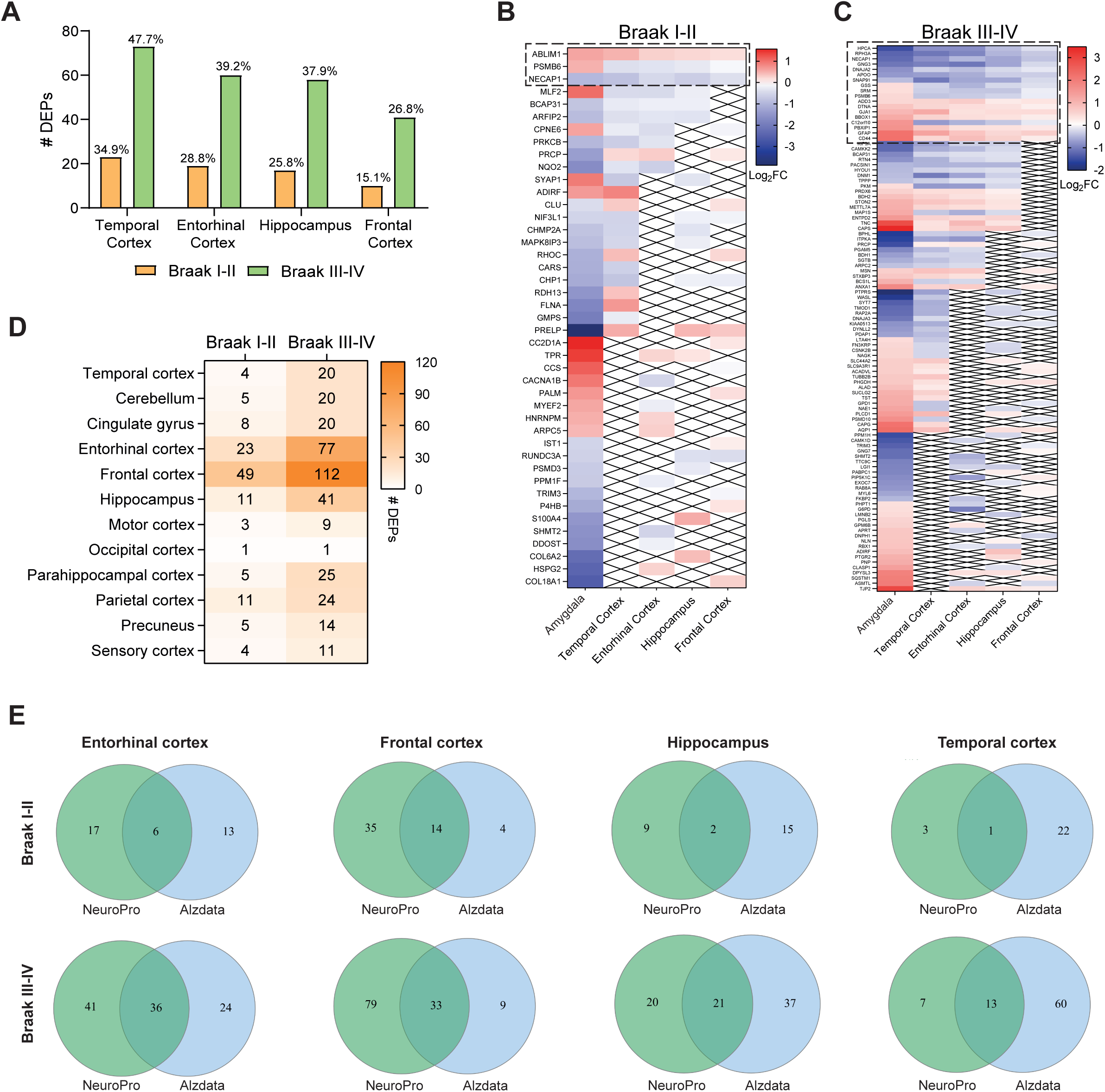
**Part of amygdaloid DEPs showed gene/protein expression correlation with AD in other brain regions**. (**A**) Percentage of proteins deregulated in Braak I-II and III-IV amygdaloid proteome that were also imbalanced at transcriptional level in other AD-affected regions published in Alzdata database [22,23]. Heatmap representing Log_2_Fold change of the proteins/genes differentially expressed in (**B**) Braak I-II or (**C**) Braak III-IV amygdala samples and other brain regions studied. (**D**) Number of common DEPs between Braak I-II or Braak III-IV amygdala proteome and 13 different brain regions from 38 published AD proteomic studies published in Neuropro database [24]. Dashed lines indicate common protein-coding genes between all brain areas studied. (**E**) Venn diagrams showing the overlap between the amygdala common DEPs deregulated in other brain regions (Neuropro) that were also altered at transcriptional level (Alzdata database). Figure created in <u>InteractiVenn.net</u> [59].

**Supplemental figure 2.**
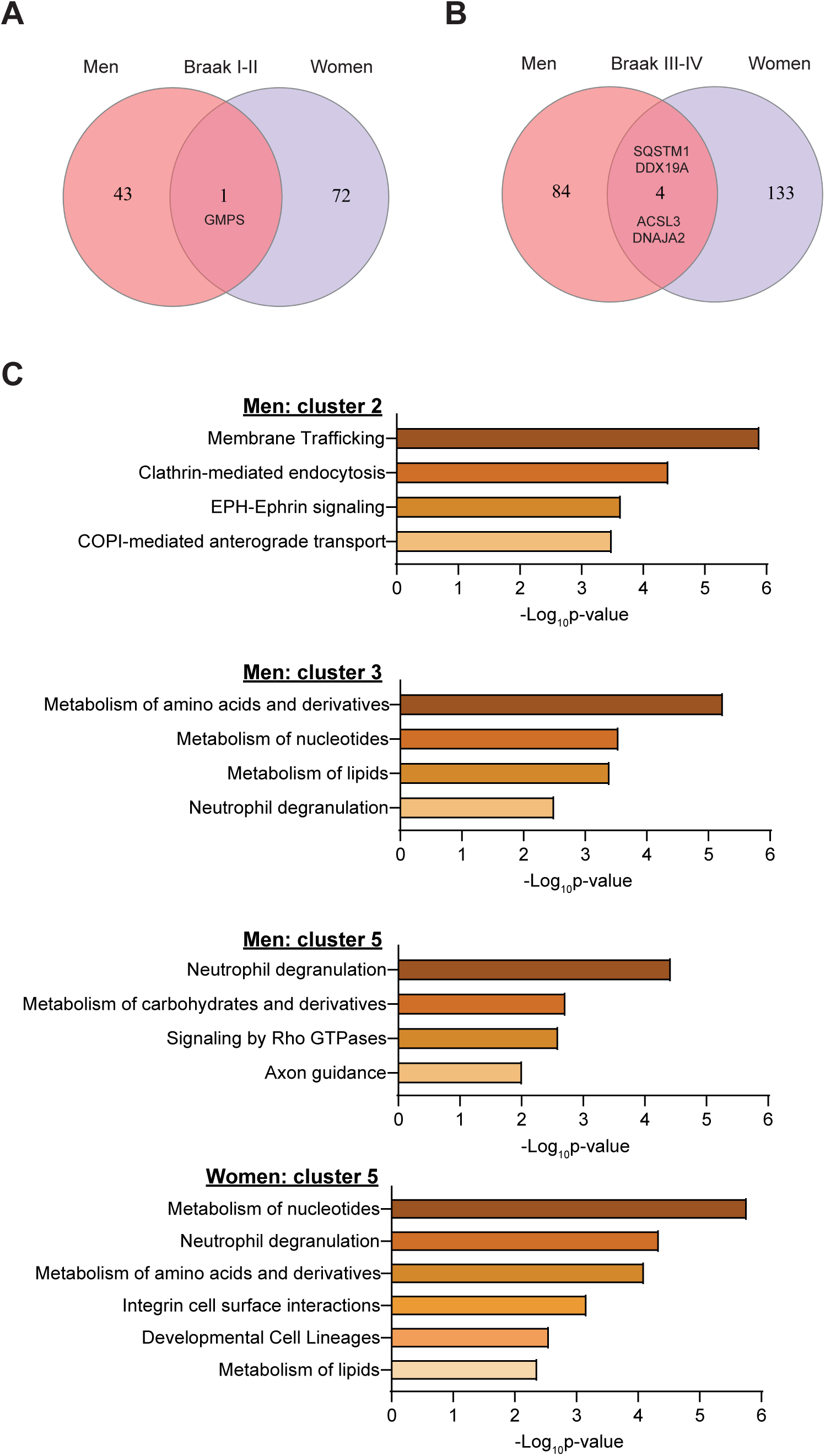
Low overlap of DEPs between women and men. Venn diagrams showing the overlap between the amygdala common DEPs deregulated in (**A**) Braak I-II and (**B**) Braak III-IV in men and women. Figure created in <u>InteractiVenn.net</u> [59]. Reactome-based pathways significantly altered in derived from the deregulated proteins of the clustering analysis.

**Supplemental Table 1.** Description of the amygdaloid samples included in this study.

**Supplemental Table 2.** Quantified proteins detected in amygdaloid samples.

**Supplemental Table 3.** Differential expressed proteins in amygdaloid Braak I-II stage compared to control group.

**Supplemental Table 4.** Differential expressed proteins in amygdaloid Braak III-IV stage compared to control group.

**Supplemental Table 5.** Common differentially expressed gene/proteins (p≤0.05) between Braak I-II or Braak III-IV amygdaloid datasets and other AD-affected brain regions from AlzData database [22,23] with their Log_2_Fold Change values.

**Supplemental Table 6.** Common differentially expressed gene/proteins (p≤0.05) between Braak I-II or Braak III-IV amygdaloid datasets and other AD-affected brain regions from Neuropro database [24]. Positive and negative values indicate protein up- or down-regulation, respectively.

**Supplemental Table 7.** Common differentially expressed gene/proteins (p≤0.05) between Braak I-II or Braak III-IV amygdaloid datasets and other AD-affected brain regions from AlzData and Neuropro databases [22–24].

**Supplemental Table 8.** Differential expressed proteins in amygdaloid men Braak I-II stage compared to control group.

**Supplemental Table 9.** Differential expressed proteins in amygdaloid men Braak III-IV stage compared to control group.

**Supplemental Table 10.** Differential expressed proteins in amygdaloid women Braak I-II stage compared to control group.

**Supplemental Table 11.** Differential expressed proteins in amygdaloid women Braak III-IV stage compared to control group.

**Supplemental Table 12**. Clustering analysis separated by sex comparing controls, Braak I-II, and Braak III-IV subjects. Statistical analyses were performed by one-way ANOVA test.

**Supplemental Table 13**. Commonly deregulated proteins (p<0.05) between Braak I-II or Braak III-IV amygdaloid datasets separated by sex and other AD-affected brain regions from Agora database. Values indicate Log2 Fold Change

**Supplemental Table 14**. Top genes whose silencing reverse the proteomic signature of amygdala DEP datasets from each Braak stage. Connectivity scores near to -100 express highly potential activity to reverse the proteomic signature. Results were obtained from connectivity map database [20].

**Supplemental Table 15**. Top genes whose silencing reverse the proteomic signature of amygdala DEP datasets from each Braak stage considering sex. Connectivity scores near to - 100 express highly potential activity to reverse the proteomic signature. Results were obtained from connectivity map database [20].

## Notes

### Competing Interest Statement

The authors have declared no competing interest.

